# Crocetin and crocin production in plant and bacteria using a CCD4 enzyme from *Nyctanthes arbor-tristis*

**DOI:** 10.1101/2025.03.13.642995

**Authors:** Lucía Morote, Elena Moreno Giménez, Alberto José López Jiménez, Ángela Rubio-Moraga, Verónica Aragonés, Oussama Ahrazem, José-Antonio Daròs, Lourdes Gómez-Gómez

**Affiliations:** Instituto Botánico. Departamento de Ciencia y Tecnología Agroforestal y Genética. Universidad de Castilla-La Mancha, Campus Universitario s/n, 02071 Albacete, Spain; Escuela Técnica Superior de Ingeniería Agronómica y de Montes y Biotecnología. Departamento de Ciencia y Tecnología Agroforestal y Genética. Universidad de Castilla-La Mancha, Campus Universitario s/n, 02071 Albacete, Spain; Facultad de Farmacia, Universidad de Castilla-La Mancha, Campus Universitario s/n, 02071 Albacete, Spain; Instituto de Biología Molecular y Celular de Plantas (Consejo Superior de Investigaciones Científicas-Universitat Politècnica de València), 46022 Valencia, Spain

**Keywords:** Aqueous two-phase, crocetin, carotenoid cleavage dioxygenase, Buddleja, Crocus, Verbascum, metabolic engineering

## Abstract

Crocins are hydrophilic crocetin esters that are arising interest as cosmetics and pharmaceuticals. Crocetin dialdehyde, the precursor of crocetin, derives from a C7-C8(C7՛-C8՛) cleavage of carotenoids in a few plant species including *Crocus sativus* or *Nyctanthes arbor-tristis*. We investigated the genome of *N. arbor-tristis* and identified an enzyme from the CCD4 subfamily catalyzing the cleavage of zeaxanthin to produce crocetin dialdehyde. This enzyme, NatCCD4.1, was used for the microbial production of crocetin dialdehyde in a two-phase culture system using direct extraction with n-dodecane, resulting in a titer of 109.2 ± 3.23 mg/L, which is the highest crocetin dialdehyde yield reported in bacteria so far. Further, a viral vector was used to express NatCCD4.1 in *Nicotiana benthamiana*, triggering a crocin accumulation of 2.32 ± 0.69 mg/g DW. Our results provide new insights into crocin biosynthesis and demonstrate that NatCCD4.1 is a valuable tool for improving crocetin and crocin production in heterologous systems.

## Introduction

Crocetin is a class of lipophilic isoprenoid molecule composed of a polyunsaturated chain, with two alcohol (crocetindiol), two aldehyde (crocetindial), two carboxylic acid (crocetin), or esters (crocins) functional groups (Figure 1) ^1^. Crocetin and crocins ^2^ display a broad range of therapeutic properties due to their antioxidant and anti-inflammatory activities and are also known to reduce the risk of certain cancers ^3, 4^. Crocetin and crocins have been tested in clinical trials on depression, anxiety, and other brain disorders as Alzheimer and Parkinson ^5–7^. These advantageous characteristics are fueling the expansion of crocetin esters in a range of pharmaceutical goods ^8^. The crocetin and crocin market size has been estimated to be about 4.7 billion USD worldwide and is expected to grow at a compound annual growth rate (CAGR) of 5.16% from 2024 to 2030. The crocetin biosynthesis pathway is present only in plants, and *Crocus sativus* stigmas and *Gardenia jasminoides* fruits are the principal commercial source of these compounds. Other plants are known to produce crocetin and crocins, but at lower levels, as *Buddleja davidii* ^9^, *Verbascum* sp. ^10^ and *Nyctanthes arbor-tristis* ^11^. The actual production of crocetin and corresponding esters cannot meet the market demand due to the high costs associated to cultivation and extraction from the main plant source ^12^. In addition, prevailing environmental conditions, such as global warming and drought, have exacerbated the production of these valuable metabolites ^13^. To meet the increasing demand for crocetin and crocins, several attempts in different heterologous systems using the enzymes from crocetin biosynthesis in saffron, *Buddleja* and gardenia have been reported with variable results ^14, 15^. New biotechnological processes for the improved production of crocetin and crocins using metabolically engineered microorganism and fast-growing plants with highly active enzymes are needed. For such purpose, discovery, and characterization of new enzymes for crocetin biosynthesis is a straightforward strategy.

**Figure 1.**
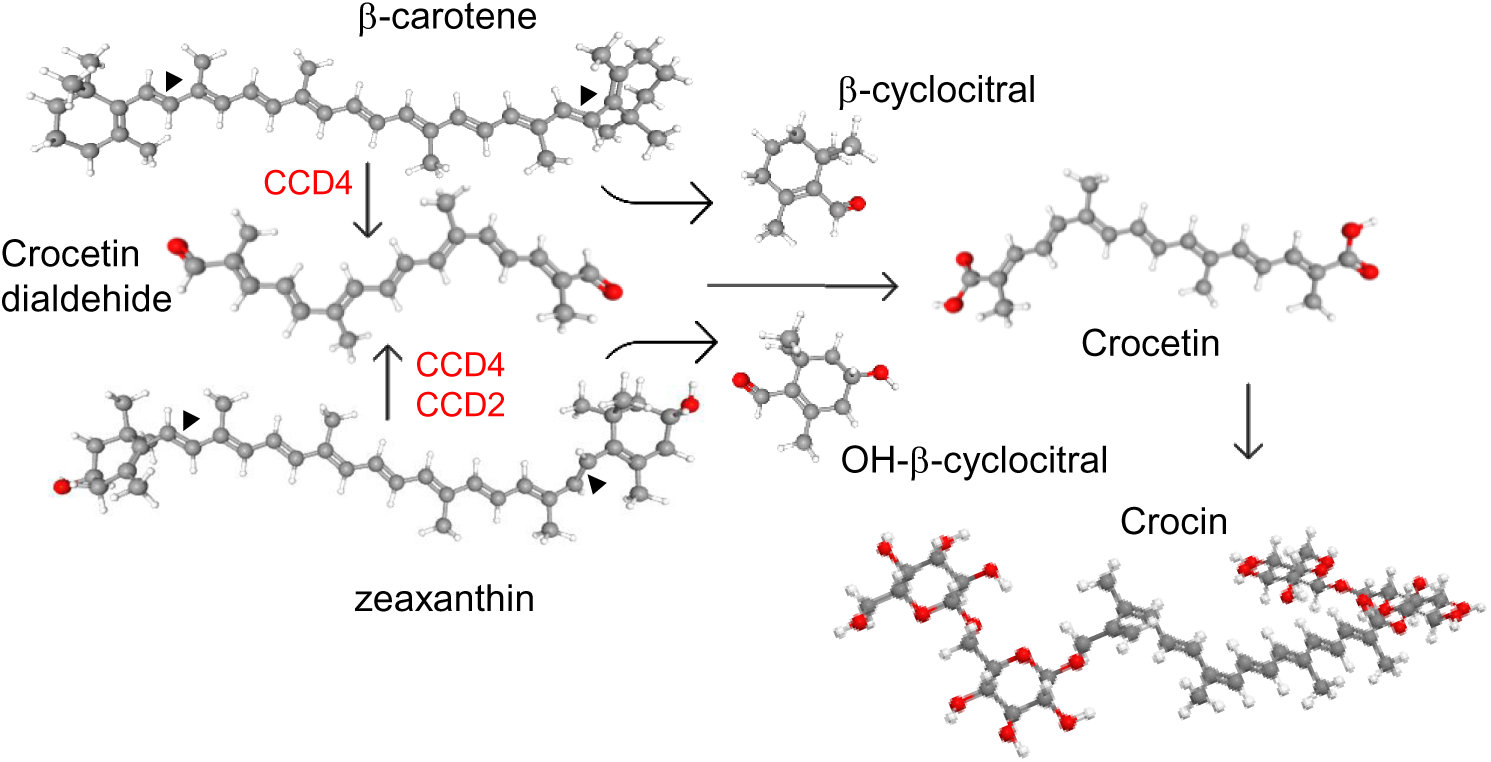
Cleavage activities of carotenoid cleavage dioxygenase (CCD) enzymes on β-carotene and zeaxanthin, producing different apocarotenoid products. In red are shown the CCD subfamilies involved in the C7-C8(C7՛-C8՛) cleavage of carotenoids.

The enzymes involved in the biosynthesis of crocetin belong to the carotenoid cleavage dioxygenase (CCD) family, which catalyze the cleavage of carotenoids to render apocarotenoid compounds ^16^. These enzymes have different substrate specificity and cleavage sites, producing a wide range of apocarotenoids ^17^. The CCD enzymes involved in the biosynthesis of crocetin cleave at the C7–C8 (C7′–C8′) double bonds of different carotenoid substrates. The enzyme CsCCD2L from *C. sativus* produces crocetin dialdehyde by cleaving zeaxanthin ^18, 19^, while in gardenia, *Buddleja* and *Verbascum*, CCDs that belong to the CCD4 subfamily catalyze the biosynthesis of crocetin dialdehyde, and can recognize one or several carotenoids as substrates, such as lycopene, zeaxanthin and β-carotene ^9, 10, 20^ (Figure 1). In addition, a CCD4 from *Bixa orellana*, which does not accumulate crocetin or crocins, has been shown to catalyzed the C7–C8 (C7′–C8′) cleavage of lycopene ^21^.

Recently, genome and transcriptome analyses of *N. arbor-tristis* have allowed the identification of 15 contigs with homologies to CCDs that could be involved in the biosynthesis of crocetin in this species ^22^. Crocins accumulate in the colored tubular calyx of the flower ^11, 23^ that also contains safranal ^24^. The total concentration of crocins in the calix reaches 35.57% (w/w) and have been used as a potential wound healing phytoconstituent ^25^, which also shows hypoglycemic and hypolipidemic properties ^26^. In this study, we have identified a CCD4 from *N. arbor-tristis* involved in the biosynthesis of crocetin. Functional analysis of this enzyme showed its C7–C8 (C7′– C8′) cleavage activity of zeaxanthin. Furthermore, to advance the biotechnological exploitation of this CCD in different heterologous platforms, a two-phase culture system using n-dodecane was used to produce crocetin dialdehyde in *Escherichia coli*, and a virus driven system to produce crocins in *Nicotiana benthamiana*.

## Results

*Identification of a CCD4 enzyme in N. arbor-tristis involved in the synthesis of crocetin.* In a previously published transcriptome of *N. arbor-tristis* tissues ^22^, 15 contigs with a high identity to CCD4-encoding genes and expressed at high levels in flower tissue were identified. From these 15 contigs, 11 of them were pedicel-specific with variable expression levels. Only two contigs out of the 15 encoded full-length CCD4 proteins. In order to get the complete sequences for all the other contigs, we analyzed the sequenced genome of *N. arbor-tristis* (Bioproject PRJEB46894). A total of 8 genes encoding CCD4 were identified and named from NatCCD4.1 to NatCCD4.8, which showed identities from 46.46% to 98.66% (Figure 2 and Supplemental Figure S1). The identified contigs in the flower transcriptome correspond to genes *NatCCD4.1*, *3*, *4*, *5* and *6*, while no contigs correspond to genes *NatCCD4.2*, *7* and *8*. All the amnio acid sequences were analyzed for predicted localization (Supplementary Figure S2), and all proteins were predicted to be targeted to the plastid.

**Figure 2.**
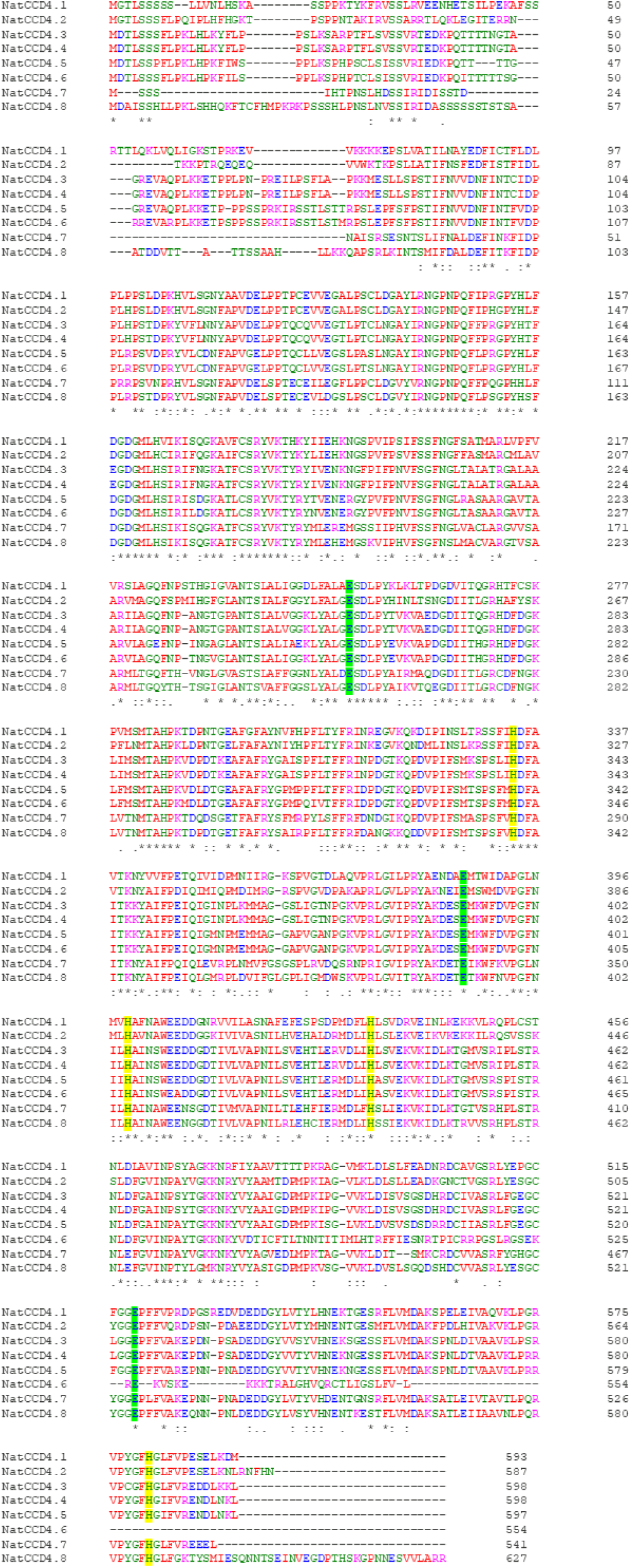
Amino acid sequence alignment of NatCCD4 enzymes. Conserved amino acid residues are depicted with asterisk. The amino acid residues involved in iron coordination are highlighted in yellow and green.

A phylogenetic tree with the identified CCDs and CCD protein sequences from *Arabidopsis thaliana* was inferred using the maximum-likelihood method (Figure 3). The CCD4 enzymes were closely related to each other, and these sequences were used to build an additional phylogenetic tree with those CCD4 sequences with characterized CCD activity in the literature (Figure 4). The CCD4 enzymes from *N. arbor-tristis* were placed in different sub-clusters. One of the subclusters containing 6 of the CCD4 enzymes identified in *N. arbor-tristis* was closely related to the cluster of CCD4 with a C9–C10 (C9′–C10′) cleavage activity, while the NatCCD4.1 and NatCCD4.2 were closely related to CCD4 enzymes from *Buddleja* and *Verbascum* which showed a C7– C8 (C7′–C8′) cleavage activity (Figure 4). This result suggested its involvement in the biosynthesis of crocins in *N. arbor-tristis*. Both genes were present in tandem in the analyzed genomic contig (NycArb205559), in the same orientation, separated by less than 2600 base pairs (bp). However, contigs for NatCCD4.2 were not present in the flower transcriptome, suggesting the lack or reduced expression of this gene in this tissue. In fact, analysis of the promoter sequences of both genes showed a 48.06% identity (Supplemental Figure S3) suggesting a differential regulation.

**Figure 3.**
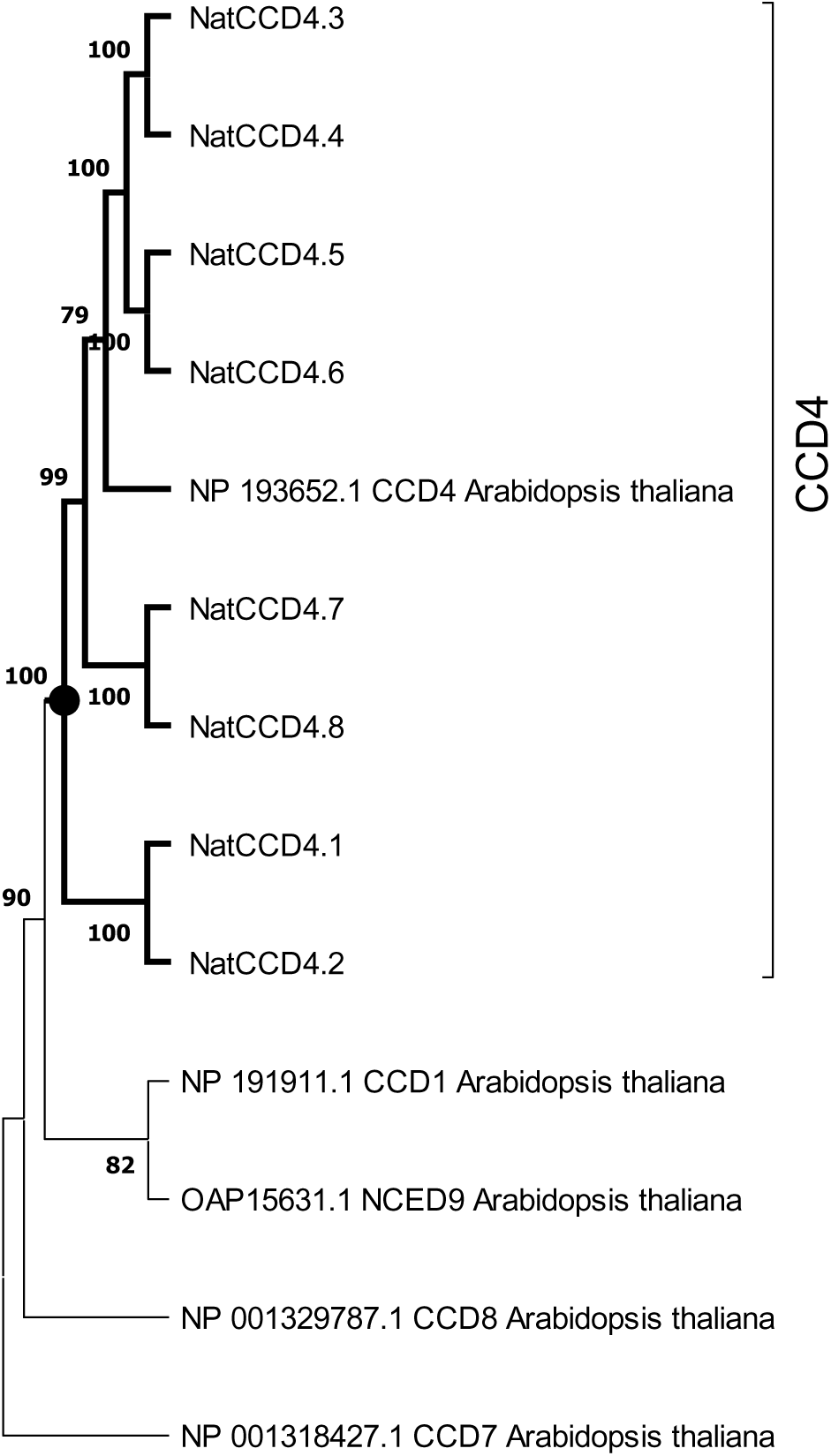
Phylogenetic tree of CCD4 encoded by genes identified in the genome of *Nyctanthes arbor-tristis*, and CCDs and NCEDs of *Arabidopsis thaliana*. The cluster containing all the CCD4 enzymes is highlight in bold. Phylogenetic analysis was done using MEGA version 11.0.10 with the maximum-likelihood method (https://megasoftware.net/) and bootstrap tests replicated 5000 times.

**Figure 4.**
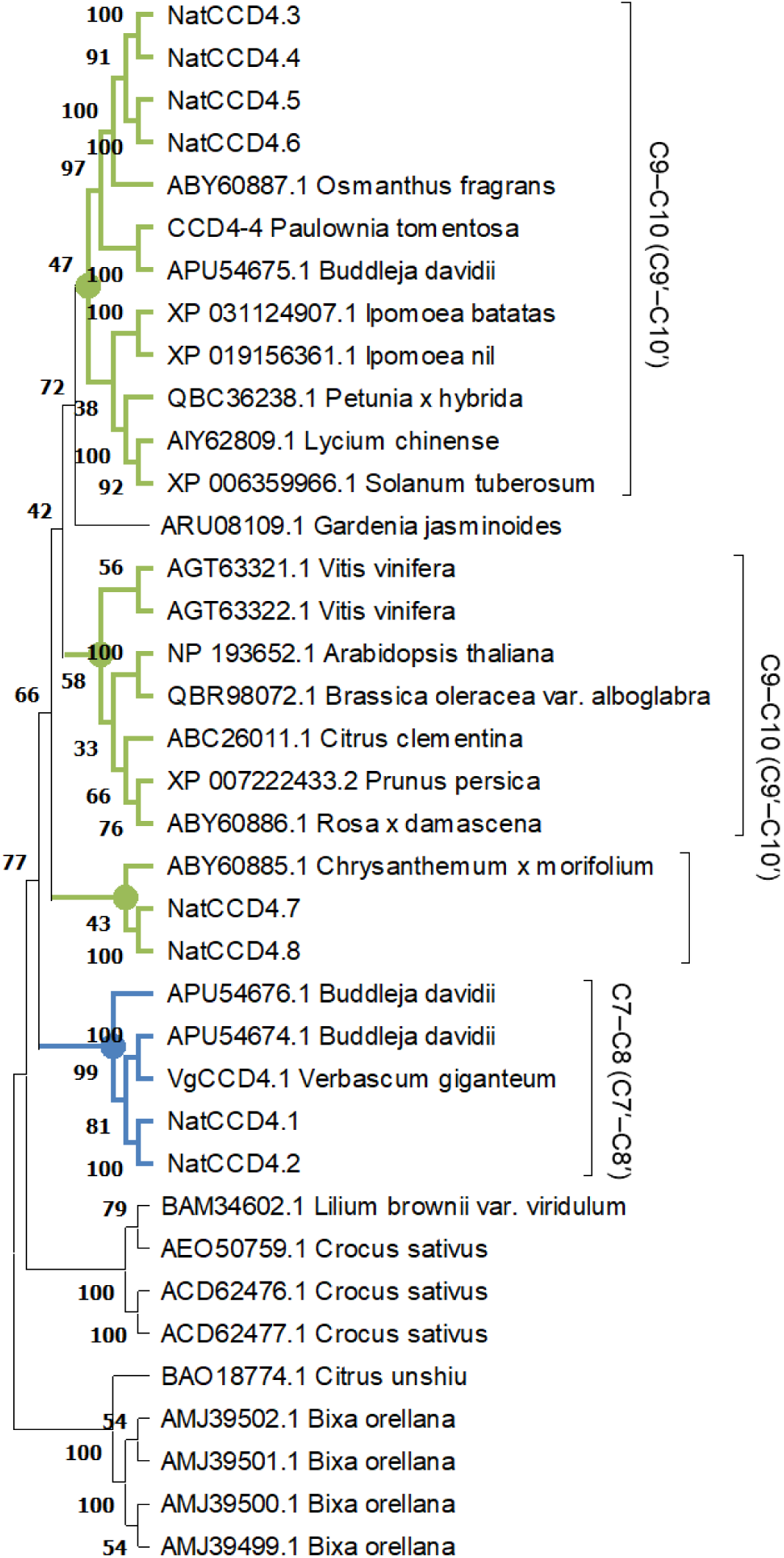
Phylogenetic relationship of CCD4 enzymes from different plant species. The phylogenetic tree was constructed by the maximum-likelihood method (https://megasoftware.net/) and bootstrap tests replicated 5000 times. The number on the nodes corresponds to the percentage of bootstrap values. The sub-tree including the CCD4 sequence with a C7-C8:C7′-C8′ cleavage activity is labelled in blue, while in green are labelled the cluster grouping enzymes which showed a C9-C10:C9′-C10′ activity. The activities reported for each CCD are shown on the right.

Both proteins, which share 68.62% identity, showed the common characteristic seven-bladed β-propeller structure (Supplementary Figure S4) with a series of loops in whose center a Fe^2+^ is localized. This cation is, essential for the catalytic activity, ensuring the activation of oxygen for cleavage of carotenoid/apocarotenoid substrates ^27^. Around this divalent cation, four conserved histidine (H) residues form the first coordinated sphere while three glutamic acid (E) residues form the second coordinated sphere (Supplementary Figure S4).

### Functional characterization of CCD4 in E. coli cells

To determine the activity of NatCCD4.1, the corresponding cDNA was cloned in the expression vector pTHIO-DAN1, affording arabinose-inducible expression. *E. coli* cells that co-express different carotenoid biosynthetic genes, allowing the synthesis of lycopene, β-carotene, or zeaxanthin, as described previously ^28^ were transformed. After induction of NatCCD4.1 expression with arabinose, carotenoids and apocarotenoids were extracted and analyzed by HPLC-DAD. Cleavage of carotenoid substrates in *E. coli* destroys the chromophore, causing a loss of color (also referred to as bleaching) (Figure 5A). The reduced levels of zeaxanthin in the cells expressing NatCCD4.1 suggested cleaving of this substrate (Figure 5A). However, under the tested conditions, the detection of the carotenoid cleavage product from culture media and organic extracts of cell pellets from bleached *E. coli* cultures was not possible, suggesting that the aldehyde cleavage products were further modified or quickly degraded in *E. coli*.

**Figure 5.**
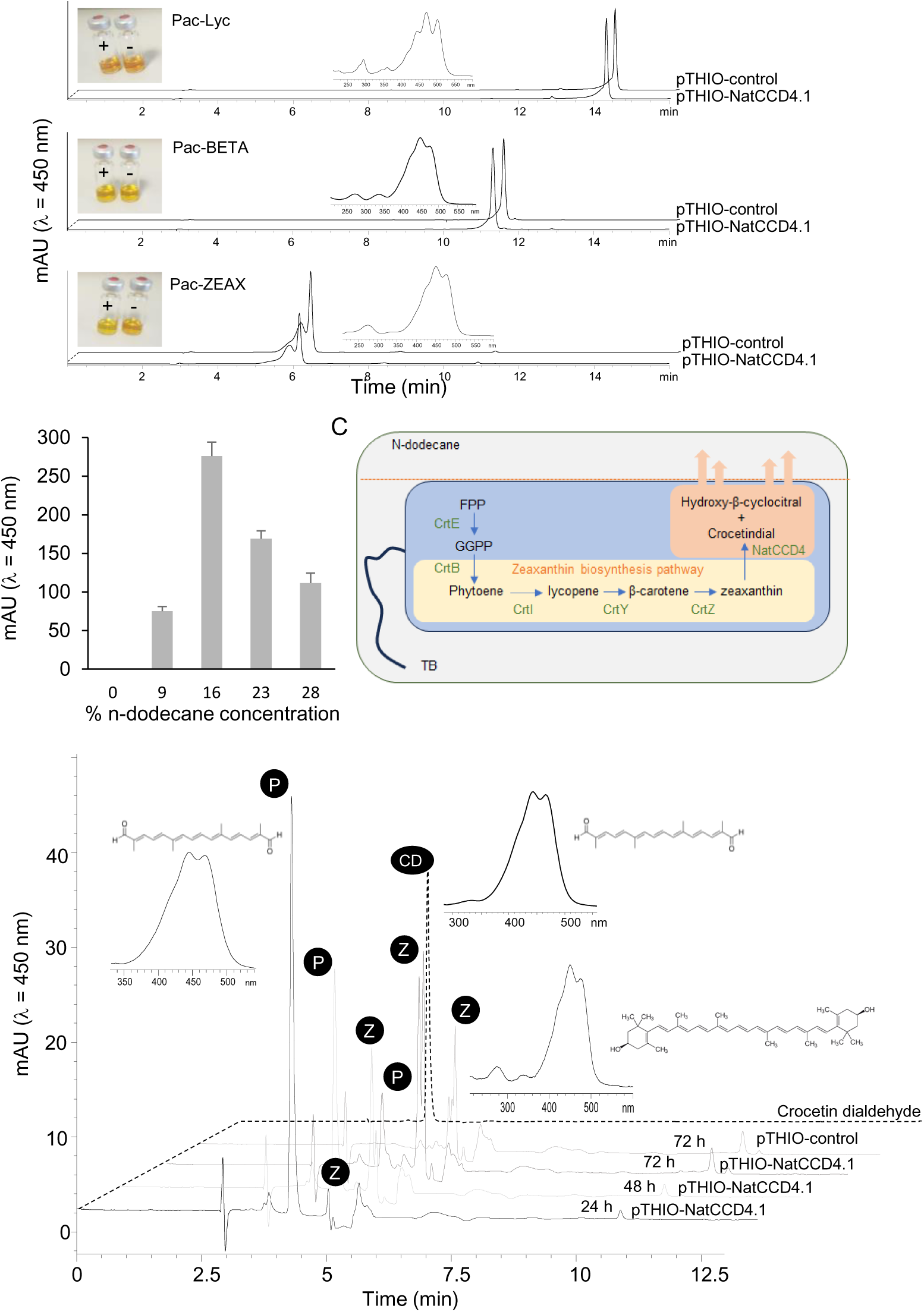
*In vivo* assays of NatCCDD4.1 using *E. coli* cells accumulating different carotenoid substrates. A) Representative HPLC-DAD chromatograms of apolar extracts obtained from *E. coli* cells expressing NatCCDD4.2 and control cells. Insets depict the UV/vis spectra of lycopene, from the extracts of cells containing the Pac-lyc plasmid, spectra of β-carotene, from the extracts of cells containing the Pac-BETA plasmid, and the spectra of zeaxanthin, from the extracts of cells containing the Pac-ZEAX plasmid. Also, the colors of the apolar extracts are shown. B) Quantification of crocetin dialdehyde obtained from *E. coli* cells producing zeaxanthin and expressing NatCCD4.1, after using different final concentrations of n-dodecane for the determination of NatCCD4.1 activity using an *in situ* two-phase culture system. C) Schematic representation of the *in situ* two-phase culture system used for the detection of NatCCD4.2 reaction product, D) Representative chromatogram profiles obtained in the HPLC-DAD analyses of NatCCD4.1 activity on zeaxanthin using the *in situ* two-phase culture system, with 16% n-dodecane final concentration and collection at different time points after arabinose induction. Insets show the structures and UV/vis spectra of crocetin dialdehyde (indicated by P) and zeaxanthin (indicated by Z).

### Two-phase culture using n-dodecane for in situ extraction of crocetin dialdehyde

To prevent apocarotenoid degradation in *E. coli*, a two-phase culture system using n-dodecane was performed for *in situ* extraction of apocarotenoids from the bacterial cells. This solvent has been previously used for the extraction of hydrophobic retinoids in *E. coli* ^29^, which significantly enhanced yield. Different final % concentrations of n-dodecane were tested to determine the optimal ratio for crocetin production. The final fractions of n-dodecane tested were 9%, 16%, 23% and 28%. The in situ extraction by n-dodecane could minimize intracellular degradation of the retinoids. Apocarotenoids were sequestered and showed to be more stable in the n-dodecane phase (Figure 5B and Figure 5C), and could not be detected in the cell mass, where the substrate zeaxanthin was retained (Supplementary Figure S5A and Figure 5C). As a result, apocarotenoid production was measured only from the n-dodecane phase. The best results were obtained with 16% final mix after 48 h of growth (Figure 5B). Further, we tested different growth times using a final 16% n-dodecane mix to determine the impact on the production of crocetin dialdehyde, from 24, 48 and 72 h. The best result was obtained after 24 h of incubation (Figure 5D), with a 109.20 ± 3.23 mg/L production of crocetin dialdehyde, compared to the 61.00 ± 1.90 mg/L and 23.60 ± 1.12 mg/L obtained after 48 and 72 h, respectively (Figure 5D). In addition, longer incubation times resulted in the extraction of more carotenoid substrate in the organic phase, which is mainly present in the cell pellet (Supplemental Figure S5B), due to cell death and lysis. The obtained results are in agreement with previous observations in *E. coli* cells expressing CsCCD2L ^30^, which showed that the decrease in crocetin dialdehyde accumulation after 27 h was related to changes in the expression level of CsCCD2 mRNA, suggesting that CsCCD2 mRNA expression/stability could be a potential target for increasing the yield of crocetin dialdehyde biosynthesis in bacteria ^30^.

Once the optimal conditions were stablished for the two-phase culture system for the *in situ* extraction of the apocarotenoids products, we decided to test the activity of NatCCD4.2 using *E. coli* cells accumulating lycopene, β-carotene, and zeaxanthin (Supplemental Figure S6). However, no products were detected using these three different carotenoids substrates under these optimized conditions.

### Functional characterization of CCD4 in N. benthamiana

Subsequently, we constructed a recombinant virus (Figure 6A) to express NatCCD4.1 (TEV-NatCCD4.1) in *N. benthamiana* plants. NatCCD4.1 was predicted to be localized in the plastid and must contain the native amino-terminal transit peptide to target the enzyme to the plastids. NatCCD4.1 was inserted in the tobacco etch virus (TEV) genome in a position that correspond to the amino terminal end of the viral polyprotein ^31^. In addition, at the 3′ end of the cloning site, the construct contains a sequence for an artificial NIaPro cleavage site to allow the release of NatCCD4.1 from the viral polyprotein. Based on our previous work, the −8/+3 NIb-CP cleavage site was used. The obtained recombinant construct TEV-NatCCD4.1 was introduced in *A. tumefaciens*, and positive clones were infiltrated in *N. benthamiana* plants. TEV-GFP that expresses GFP between viral NIb and CP cistrons was used as a control. Symptoms of infection were observed approximately 8 dpi in all plants agroinoculated with TEV-NatCCD4.1 and TEV-GFP. A unique yellow pigmentation was observed in tissues of plants agroinoculated with TEV-NatCCD4.1 at approximately 14 dpi (Figure 6B). Symptomatic leaves from plants infected with TEV-GFP, and TEV-NatCCD4.1 were collected and subjected to extraction and analysis to determine their crocin and carotenoid profiles. Analysis of the polar fraction of leaves infected with TEV-NatCCD4.1 showed a series of peaks with maximum absorbance around 440 nm, corresponding to crocins with different degree of glucosylation (Figure 6C). These peaks were absent in the extracts from tissues from mock-inoculated control plants or plants infected with TEV-GFP. Subsequently, the levels of carotenoids and chlorophylls in the apolar fractions were also investigated (Supplementary Figure S7). The comparison of the profiles of apolar extracts from tissues infected with TEV-GFP and TEV-NatCCD4.1 showed reduced levels of chlorophyll and pheophytin in TEV-NatCCD4.1. At the carotenoid level, major differences were observed in the level of zeaxanthin, with a 96.61% reduction, followed by an 87.71% reduction of β-carotene, and 75.70% reduction of lutein levels in the leaves of TEV-NatCCD4.1-infected plants (Supplementary Figure S7), which further reinforced the additional activity of this enzyme in targeting zeaxanthin. Overall, these results revealed a remarkable accumulation of 2.32 ± 0.69 mg/g of crocins in *N. benthamiana* dry weight (DW) leaf tissue.

**Figure 6.**
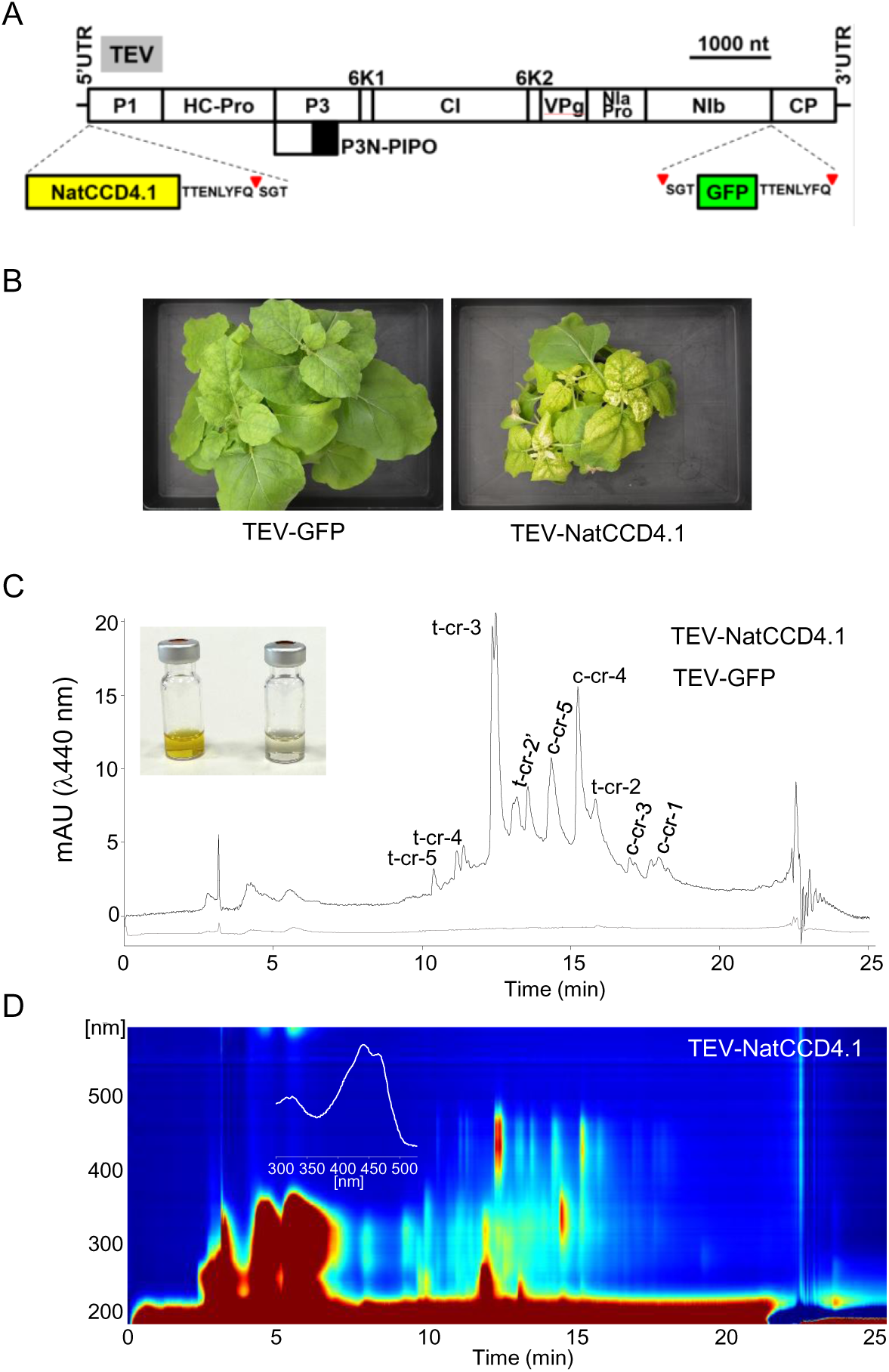
Crocin production in *N. benthamiana* plants using a TEV recombinant clone that expresses NatCCD4.1. A) Schematic representation of TEV genome indicating the position where the GFP (green box) or NatCCD4.1 (yellow box) were inserted. The sequence of the artificial NIaPro cleavage sites to mediate the release of the recombinant proteins from the viral polyprotein is indicated. The red arrow points the cleavage position. Lines represent TEV 5’ and 3’ UTR and boxes represent P1, HC-Pro, P3, P3N-PIPO, 6K1, CI, 6K2, VPg, NIaPro, NIb and CP cistrons, as indicated. Scale bar corresponds to 1000 nt. B) Pictures of representative leaves from plants mock-inoculated and agroinoculated with TEV-GFP and TEV-NatCCD4.1, as indicated, taken at 14 dpi. Scale bars correspond to 5 mm. C) Representative chromatographic profile run on an HPLC-DAD/UV system and detected at 440 nm of the polar extracts of tissues infected with TEV-GFP, and TEV-NatCCD4.1. Peak abbreviations correspond to: c-cr, cis-crocetin; t-cr, trans-crocetin; c-cr1, cis-crocin 1; t-cr1, trans-crocin 1; t-cr2, trans-crocin 2; t-cr2ʹ, trans-crocin 2ʹ; c-cr3, cis-crocin 3, t-cr3, trans-crocin 3; c-cr4, cis-crocin 4; t-cr4, trans-crocin 4; c-cr5, cis-crocin 5; t-cr5, trans-crocetin 5. D) HPLC-DAD/UV isoplot chromatogram of polar extracts and absorbance spectra of the major crocin detected in the polar extracts of *N. benthamiana* tissues infected with TEVΔNIb-NatCCD4.1. Analyses were performed at 13 dpi.

## Discussion

Crocins have a wide range of biological properties and represent valuable metabolites with multiple applications in different industrial sectors as pharmaceuticals, nutrients, and cosmetic ingredients ^3, 8^. Current commercial production of crocins is based on purification from saffron stigmas and gardenia fruits. In addition, crocin biosynthesis has been evaluated in bacteria ^30, 32^, yeast ^33^, and plants using the CCD enzymes from saffron, gardenia, *Bixa*, *Buddleja,* and *Verbascum* ^10, 14^. Here, we report the production of crocetin in *E. coli* and crocins in *N. benthamiana* using a CCD4 enzyme identified in *N. arbor-tristis*. The analysis of the genome of *N. arbor-tristis* allowed the identification of two CCD4 genes, which shared 68.62% identity, encoding for CCD4 enzymes that cluster with previous CCD4 enzymes involved in crocetin biosynthesis in *Verbascum* ^10^ and *Buddleja* ^9^. However, only one of these two proteins, NatCCD4.1, was active for crocetin dialdehyde biosynthesis. All these three plants, *N. arbor-tristis*, *Buddleja*, and *Verbascum*, belong to the Scrophulariaceae family, which suggests that the specialization of these enzymes to produce crocetin dialdehyde occurred before the divergence of these three species. As for the case of the enzymes characterized in *Verbascum* and *Buddleja,* NatCCD4.1 showed high specificity for zeaxanthin, which is in agreement with the detection of safranal among the volatiles emitted by the flowers of *N. arbor-tristis* ^24, 34^, *Buddleja* and *Verbascum* ^10, 35^. Although, the enzyme from *Verbascum* also recognizes β-carotene as substrate ^10^. In *Verbascum,* crocins are homogenously distributed in the petals of the flower, which showed a yellow coloration, while in *N. arbor-tristis* and *Buddleja* the accumulation of crocins is specifically restricted to the calix (Supplemental Figure S8), which can serve as visual cues for pollinators, enhancing the success of plant pollination.

Apocarotenoids are, in general, highly sensitive to light, temperature, and oxygen ^36^, and crocetin is not an exception ^37, 38^. Therefore, apocarotenoids produced in heterologous microbial hosts are often vulnerable to degradation ^39^. To reduce the intracellular degradation of crocetin dialdehyde, and to solve the problem of limited solubility, a two-phase *in situ* extraction using n-dodecane has been applied for retinoid production in *E. coli* and *S. cerevisiae* ^40, 41^. This strategy was used in this study to improve the detection of crocetin dialdehyde in the bacterial assay. By using this methodology, crocetin dialdehyde was recovered in the organic phase 24 h after the induction of NatCCD4.1 expression in the bacterial cultures, with low levels of zeaxanthin detected in this phase. Therefore, this methodology allowed efficient recovery of crocetin-dialdehyde without the necessity of additional high-cost processing steps such a sequentially cell-disruption and purification of the pigments from the cell pellet using organic solvents. In addition, the obtained yield of crocetin dialdehyde was higher than those levels previously reported in *E. coli* using the CsCCD2L enzyme from saffron, 5.14 mg/L ^30^ and 4.24 mg/L ^42^, and also above of the levels obtained using yeast; in this case the expression of CsCCD2L in this host produced 1.21 mg/L of crocetin ^43^. Further optimization of the system allowed the obtention of 1.95 mg/L of crocetin ^44^. In another strategy developed in yeast, using a temperature-responsive crocetin-producing strain and increasing the copy number of *CsCCD2L* gene using the CRISPR-Cas9 based multiplex genome integration technology, 1.05 mg/L were produced, suggesting the existence of different bottleneck points for crocetin production in yeast ^33^. In any case, the industrial potential of *E. coli* as a sustainable platform for crocetin-dialdehyde production is endorsed by the inexpensive starting material, allowing a one-step process that has significant advantages compared with the currently used methodology for the extraction of crocetin-dialdehyde from saffron or gardenia using different chemicals and processes needed to convert crocins into crocetin dialdehyde ^3^. In addition, the rapid growth and low growth requirements contribute to the fast production of this metabolite at reduced cost.

The transient expression using *N. benthamiana* has become the preferred plant-based platform due to its advantage in metabolites production yield and speed, as well as the lack of concern for transgene escape and contamination of food crops. Virus-drive expression of NatCCD4.1 in *N. benthamiana* plants allowed the accumulation of remarkable amounts of 2.32 ± 0.69 mg of crocins per gram (DW) of infected tissues in only 14 dpi. Previous experiments using this system and different CCDs from species accumulating crocins, showed variable levels of crocins accumulation in leaves. The CCDs previously tested include BdCCD4.1, CsCCD2L, and VgCCD4.1 ^10, 31^. Remarkably, all are in the range of mg/g of DW tissues production of crocins in *N. benthamiana* leaves. These levels are easily reached by expressing just a single gene, which highlights the advantage of plants over other hosts, such as yeasts or bacteria, which need the introduction of additional genes to biosynthesize the carotenoid substrates, aldehyde dehydrogenase and glucosyltransferases. Plants possess the basic precursors necessary to produce crocins, and additional modifier enzymes that guarantee the stability and storage of these compounds.

### Limitations of the study

Significant levels of crocetin dialdehyde and crocins have been produced in this study by the expression of a single identified gene in *N. arbor-tristis*, NatCCD4.1, in bacteria and in planta, respectively. The obtained levels are in the order of mg/L and mg/g (DW), without the introduction of further optimization procedures that can improve the current crocetin titer in *E. coli* and the crocins content in *N. benthamiana*. Improvements could be obtained from development of fermentation processes (such as fed-batch cultivation), or media formulations, in the case of crocetin dialdehyde production in *E. coli*, or the introduction of additional genes that could enhance the levels of precursor in the case of crocins biosynthesis in *N. benthamiana*.

## Acknowledgements

This work was supported by grants PID2020-114761RB and PID2020-114691RB-I00 from the Ministerio de Ciencia, Innovación y Universidades (Spain) through the Agencia Estatal de Investigación (MCIN/AEI/10.13039/501100011033). LGG is a participant of the CARNET network (RED2022-134577-T).

## Author contributions

Conceptualization: O.A. and L.G-G.; writing, reviewing, and editing: L.G-G., O.A., J-A.D. and A.R-M.; plasmids construction, E.M.J, L.M. and V.A.; plant agroinfiltration, L.M., and V.A.; metabolite extraction and analyses, L.G-G., A.R-M., and L.M.; activity assays, L.M., L.G-G., and E.M.G. Bioinformatics analyses A.J.L.J. All the authors read and approved the manuscript.

## Competing interests

The authors declare that they have no competing interests.

## Funding

This work was supported by grants PID2020-114761RB-I00 and PID2020-114691RB-I00 from the Ministerio de Ciencia, Innovación y Universidades (Spain) through the Agencia Estatal de Investigación (MCIN/AEI /10.13039/501100011033). LGG is a participant of the CARNET network (RED2022-134577-T) from the MCIN/AEI.

## Supplemental information

Supplemental Figures: S1-S8.

## STAR★Methods

### Resource availability

#### Lead contact

Further information and requests for resources and reagents should be directed to and will be fulfilled by the Lead Contact, Dr Lourdes Gómez Gómez.

#### Materials availability

This study did not generate new unique reagents and the materials generated in this study are available from the corresponding author with a completed Materials Transfer Agreement.

This study did not generate new unique reagents.

#### Data and code availability

This study did not generate standardized datasets.

This study did not report any original code.

Any additional information required to reanalyze the data reported in this paper is available from the lead contact upon request.

Experimental model and subject details

#### Plant materials and growth conditions

##### Plant growth

*Nicotiana benthamiana* was used in this study for activity assays. *N. benthamiana* seeds were sown in soil and after 10 days individual seedlings were transplanted into 12 cm diameter pots. Plants were grown under controlled environmental conditions at 25°C with a 16-h light period (light intensity 10 klux). Plants were used for infiltration 3-4 weeks after transplanting.

##### Bacterial growth

For leaf infiltration of *N. benthamiana* plants, constructs were transformed into *Agrobacterium tumefaciens* C58C1 competent cells, containing the helper plasmid pCLEAN-S48, by electroporation. Cultures were grown at 28°C with shaking overnight in Luria-Bertani (LB) liquid medium containing 50 µg/mL kanamycin, 50 µg/mL rifampicin, and 7.5 µg/mL tetracycline. Cells were collected by centrifugation and resuspended in 10 mM 2-(N-morpholino)ethanesulfonic acid (MES)-NaOH, 10 mM MgCl_2_, 150 μM acetosyringone, pH 5.6. Bacterial suspensions were adjusted at the indicated 0.5 optical density (600 nm) and infiltrated in the abaxial side of one leaf of 4-week-old plants using 1-ml needle-less syringes.

*Escherichia coli* strain BL21 (DE3) was the host strain used for the activity assays using different carotenoid substrates. Chemically competent cells were transformed with the plasmids PAC-LYC, PAC-ZEAX, and PAC-BETA (https://www.addgene.org) to produce lycopene, zeaxanthin, and β-carotene, and the positive transformants selected on LB plates containing chloramphenicol (60 µg/mL). The positive transformed bacteria were prepared for transformation by electroporation and further used for the introduction of pTHIO-NatCCD4.1, pTHIO-NatCCD4.2 and the empty vector pTHIO-Dan1. Positive double transformant cells were selected on LB plates containing ampicillin (100 µg/mL) and chloramphenicol (60 µg/mL). Double transformants were cultured overnight at 37 °C in 3 mL LB medium supplemented with ampicillin (100 µg/mL) and chloramphenicol (60 µg/mL).

#### Method details

##### Identification of CCD4 genes by bioinformatic analyses

A transcriptome from flowers of *Nyctanthes arbor-tristis* and the genome sequence [20] were searched for contigs bearing homology to genes encoding for enzymes of the CCD4 subfamily using BLAST (https://blast.ncbi.nlm.nih.gov/Blast.cgi). Phylogenetic trees were generated using MEGA version 11.0.10 with the maximum-likelihood method (https://megasoftware.net/) and bootstrap tests replicated 5000 times. Prediction of subcellular localization was obtained using the DeepLoc-1.0 software (https://services.healthtech.dtu.dk/services/DeepLoc-1.0/). The 3D structures were predicted using the Phyre2 software at intensive mode (http://www.sbg.bio.ic.ac.uk/phyre2/), and Chimera X (https://www.cgl.ucsf.edu/chimerax/).

##### Gene synthesis and cloning

DNA sequences were synthesized by the gene synthesis service of NZYtec (https://www.nzytech.com/), and used as templates for amplification with specific primers (Supplementary Table S1) for cloning in the expression vector pTHIO-Dan1 using In-Fusion assembly (Takara Bio, Europe), and for cloning in a viral vector previously described ^31^, derived from tobacco etch virus (TEV) by Gibson DNA assembly (New England Biolabs). The obtained plasmids, pTHIO-NatCCD4.1, pTHIO-NatCCD4.2 and pTEV-NatCCD4.1, were prepared with the Minitools Miniprep Kit (Biotools B&M Labs S.A., Madrid, Spain) and sequenced using an automated DNA sequencer (ABI PRISM 3730xl, Perkin Elmer, Macrogen Inc., Seoul, Korea).

##### Analysis of enzymatic activity in *E. coli*

*E. coli* strain BL21 (DE3) was the host strain used for the activity assays using different carotenoid substrates. Purchased chemically competent cells were transformed with the plasmids PAC-LYC, PAC-ZEAX, and PAC-BETA (https://www.addgene.org) to produce lycopene, zeaxanthin, and β-carotene. The positive transformants were grow in LB supplemented with chloramphenicol (60 µg/mL) and prepared for electroporation using a Micropulser Electroporator (BioRad, Hercules, Calif.) with a field strength of 18 kV/cm, for the introduction of pTHIO-NatCCD4.1, pTHIO-NatCCD4.2 and the empty vector pTHIO-Dan1. Positive electro transformed clones were selected in LB plates supplemented with ampicillin (100 µg/mL) and chloramphenicol (60 µg/mL). Double transformants were cultured overnight at 37 °C in 3 mL LB medium supplemented with ampicillin (100 µg/mL) and chloramphenicol (60 µg/mL). Cultures for activity assays were carried out in 50 mL of 2x YT (16 g of tryptone, 10 g of yeast extract, 5 g of NaCl) or in Terrific Broth (24 g of yeast extract, 20 g of tryptone, and 4 mL of glycerol per liter, and 0.017 M KH2PO4, and 0.072 M K2HPO4) supplemented with ampicillin (50 µg/mL) and chloramphenicol (30 µg/mL) using a shaking incubator at 30°C and 200 revolution per min (rpm). Expression was induced with 2% (w/v) arabinose when optical density at 600 nm (OD600) of 0.6 was reached. For the two-phase culture for apocarotenoid production, n-dodecane was layered over the culture medium (Terrific Broth) immediately after arabinose induction using different final concentrations (9%, 16%, 23% and 28%) and incubation times (6, 24, 36 and 72 h).

##### Analysis of carotenoids and apocarotenoids

Carotenoid and apocarotenoid products were extracted from bacterial cell pellets as previously described ^28^. Basically, the cells were collected by centrifugation for 10 min at 12,000 x g, the medium was separated, and acetone was added to the cell pellets, followed by the addition of an equal volume of MeOH:CHCl_3_ (1:1), mixed by vortex and sonicated in a water-bath for 10 min. The extract was centrifuge at 9,000g for 10 min and the liquid collect and evaporated for further HPLC analyses. In the two-phase culture system with a n-dodecane overlay, the upper organic phase containing the CCD cleavage products was collected and centrifuged for 10 min at 12,000 x g to remove cellular remains and directly used for apocarotenoid analyses. The pigments extracted from the cell pellets and those present in the n-dodecane phases were analyzed by HPLC-DAD (Agilent technologies 1100 series) at detection wavelengths of 450 nm and using a YMC C30 (250 x 4.6 mm, 5µm) column (Waters, Milford, USA). The mobile phases were 98:2 methanol (A), 95:5 methanol (B) and 100% methyl tert-butyl ether. The column was developed at a flow rate of 1 mL min−1 with the following gradient elution: 80% A, 20% C at 0 min, followed by linear gradient to 60% A, 40% C to 3 min at 4 min with gradient changing to 60% B, 40% C followed by a linear gradient to 0% B, 100% C by 12 min and return to initial conditions by 13 min. A re-equilibration (10 min) was carried out at initial conditions of 80% A, 20% C. A flow rate of 1 mL/min and column temperature of 40°C were used. Crocetin dialdehyde (Cat. No. 18804, Sigma) was used as standard compound. The results are presented as means ± standard deviation (SD) from three independent experiments.

##### Activity assays in *N. benthamiana*

The viral recombinant clone TEV-NatCCD4.1 consists of a wild-type TEV cDNA (GenBank DQ986288, containing the silent and neutral mutations G273A and A1119G), in which the NatCCD4.1 cDNA was inserted as the amino terminal cistron and was followed with an artificial NIaPro cleavage site to release the enzyme from the viral polyprotein. This viral clone was flanked by the 35S promoter and terminator sequences from the cauliflower mosaic virus (CaMV), in a binary vector derived from pCLEAN-G181 ^45^. The control construct TEV-GFP contains the cDNA of the enhanced GFP between NIb and CP cistrons.

*Agrobacterium tumefaciens* C58C1 competent cells, containing the helper plasmid pCLEAN-S48, were transformed by electroporation with pGTEV-NatCCD4.1 or pGTEV-GFP, as control. Transformants were selected in plates with 50 µg/mL kanamycin, 50 µg/mL rifampicin, and 7.5 µg/mL tetracycline. Selected clones were grown in liquid media and were prepared to infiltrate two leaves of one-month-old *N. benthamiana*. Inoculated plants were kept under controlled conditions in a growth chamber at 25°C under a 16/8 h day-night photoperiod. Leaf tissue was collected 14 days post-inoculation (dpi), frozen immediately, and lyophilized for further analysis for apocarotenoid and carotenoid content.

##### Extraction and analysis of apocarotenoids and carotenoids from *N. benthamiana* leaves by TLC and HPLC-DAD

Lyophilized tissues were ground with a mixer mill MM400 (Retsch GmbH, Haan, Germany) in 2 mL tubes and 0.05 g used for polar and apolar extractions. For extraction of crocins, 50% methanol was added to the homogenate, mixed, vortexed, sonicated during 10 min in a water bath, and centrifuged 10 min at 10.000 x g. The supernatant was retained for crocins analyses by HPLC-DAD by suing a method previously described ^10^. In brief, 20 μL were injected to determine its crocin content using HPLC (Konik HPLC system (Barcelona, Spain) with a DAD wavelength detector set to 440 nm and a C18 column (250×4.6 mm, 5 µM). The column was kept at 30 °C. Mobile phase A (10% acetonitrile in water and 0.01% TFA) B (90% acetonitrile and 0.01% TFA), and mobile phase C (100% methanol) were used for gradient elution at 0.8 mL/min as follows: 82.5% A and 12.5% B for 5 min, 18.8% A and 81.2% B for 15 min, and 100% C for 15 min. Further, the obtained pellet was extracted with 2:1 methanol:chloroform, mixed for an additional 10 min and incubated in an ultrasound bath for an additional 10 min followed by centrifugation. The apolar phase containing carotenoids and chlorophylls was evaporated under N_2_ gas and the dried pellet was stored with the polar extracts at −80 °C until analysis by HPLC-DAD. All assays were performed in triplicate. The HPLC-DAD method used for the analysis and detection of carotenoids was the same described in the bacteria assay section. Metabolite identification was done by comparison of retention times, and UV-visible spectra using zeaxanthin, β-carotene, lycopene and lutein standards purchased from CaroteNature (Lupsingen, Switzerland).

Thin layer chromatography (TLC) for carotenoid and chlorophyll separation was performed with silica gel 60 F254 plates using petroleum ether: diethyl ether: acetone (30:15:15) as the mobile phase.

## Notes

### Competing Interest Statement

The authors have declared no competing interest.

