## Supplementary material for "Crocetin and crocin production in plant and bacteria using a CCD4 enzyme from *Nyctanthes arbor-tristis*": FIGURES SUP.

Fig. S1

```
>NatCCD4.1
MTSMGTLSSSSSLLVNLHSKASSPPKTYKFRVSSLRVEENHETSILPEKAFSSRTTLQKLVQLIGKSTPRKEVVKKKKEPSLVATILNAYEDFICTFLDL
PLPPSLDPKHVLSGNYAAVDELPPTPCEVVEGALPSCLDGAYLRNGPNPQFIPRGPYHLFDGDGMLHVIKISQGKAVFCSRYVKTHKYIIEHKNGSPVIP
SIFSSFNGFSATMARLVPFVVRSLAGQFNPSTHGIGVANTSLALIGGDLFALAESDLPYKLKLTPDGDVITQGRHTFCSKPVMSMTAHPKTDPNTGEAFG
FAYNVFHPFLTYFRINREGVKQKDIPINSLTRSSFIHDFAVTKNYVVFPETQIVIDPMNIIRGKSPVGTDLAQVPRLGILPRYAENDAEMTWIDAPGLNM
VHAFNAWEEDDGNRVVILASNAFEFESPSDPMDFLHLSVDRVEINLKEKKVLRQPLCSTNLDLAVINPSYAGKKNRFIYAAVTTTTPKRAGVMKLDLSLF
EADNRDCAVGSRLYEPGCFGGEPFFVPRDPGSREDVDEDDGYLVTYLHNEKTGESRFLVMDAKSPELEIVAQVKLPGRVPYGFHGLFVPESELKDM
>NatCCD4.2
MGTLSSSFLPQIPLHFHGKTPSPPNTAKIRVSSARRTLQKLEGITERRNTKKPTRQEQEQVVWKTKPSLLATIFNSFEDFISTFIDLPLHPSLDPKHVLS
GNFAPVDELPPTPCEVVEGALPSCLDGAYIRNGPNPQFIPHGPYHLFDGDGMLHCIRIFQGKAIFCSRYVKTYKYLIEHKNGSPVFPSVFSSFNGFFASM
ARCMLAVARVMAGQFSPMIHGFGLANTSIALFGGYLFALGESDLPYHINLTSNGDIITLGRHAFYSKPFLNMTAHPKLDPNTGELFAFAYNIYHPFLTYF
RINKEGVKQNDMLINSLKRSSFIHDFAVTKNYAIFPDIQIMIQPMDIMRGRSPVGVDPAKAPRLGVLPRYAKNEIEMSWMDVPGFNMLHAVNAWEEDDGG
KIVIVASNILHVEHALDRMDLIHLSLEKVEIKVKEKKILRQSVSSKSLDFGVINPAYVGKKNRYVYAAMTDPMPKIAGVLKLDLSLLEADKGNCTVGSRL
YESGCYGGEPFFVQRDPSNPDAEEDDGYLVTYMHNENTGESMFLVMDAKFPDLHIVAKVKLPGRVPYGFHGLFVPESELKNLRNFHN
>NatCCD4.3
MDTLSSSFLPKLHLKYFLPPSLKSARPTFLSVSSVRTEDKPQTTTTNGTAGREVAQPLKKETPPLPNPREILPSFLAPKKMESLLSPSTIFNVVDNFINT
CIDPPLHPSTDPKYVFLNNYAPVDELPPTQCQVVEGTLPTCLNGAYIRNGPNPQFFPRGPYHTFEGDGMLHSIRIFNGKATFCSRYVKTYRYIVENKNGF
PIFPNVFSGFNGLTALATRGALAAARILAGQFNPANGTGPANTSLALVGGKLYALGESDLPYTVKVAEDGDIITQGRHDFDGKLIMSMTAHPKVDPDTKE
AFAFRYGAISPFLTFFRINPDGTKQPDVPIFSMKSPSLIHDFAITKKYAIFPEIQIGINPLKMMAGGSLIGTNPGKVPRLGVIPRYAKDESEMKWFDVPG
FNILHAINSWEEDDGDTIVLVAPNILSVEHTLERVDLIHLSVEKVKIDLKTGMVSRIPLSTRNLDFGAINPSYTGKKNKYVYAAIGDPMPKIPGVVKLDI
SVSGSDHRDCIVASRLFGEGCLGGEPFFVAKEPDNPSADEDDGYVVSYVHNEKSGESSFLVMDAKSPNLDIVAAVKLPSRVPCGFHGLFVREDDLKKL
>NatCCD4.4
MDTLSSSFLPKLHLKYFLPPSLKSARPTFLSVSSVRTEDKPQTTTTNGTAGREVAQPLKKETPPLPNPREILPSFLAPKKMESLLSPSTIFNVVDNFINT
CIDPPLHPSTDPKYVFLNNYAPVDELPPTQCQVVEGTLPTCLNGAYIRNGPNPQFFPRGPYHTFEGDGMLHSIRIFNGKATFCSRYVKTYRYIVENKNGF
PIFPNVFSGFNGLTALATRGALAAARILAGQFNPANGTGPANTSLALVGGKLYALGESDLPYTVKVAEDGDIITQGRHDFDGKLIMSMTAHPKVDPDTKE
AFAFRYGAISPFLTFFRINPDGTKQPDVPIFSMKSPSLIHDFAITKKYAIFPEIQIGINPLKMMAGGSLIGTNPGKVPRLGVIPRYAKDESEMKWFDVPG
FNILHAINSWEEDDGDTIVLVAPNILSVEHTLERVDLIHLSVEKVKIDLKTGMVSRIPLSTRNLDFGAINPSYTGKKNKYVYAAIGDPMPKIPGVVKLDI
SVSGSDHRDCIVASRLFGEGCLGGEPFFVAKEPDNPSADEDDGYVVTYVHNEKNGESSFLVMDAKSPNLDTVAAVKLPRRVPYGFHGIFVRENDLNKL
>NatCCD4.5
MDTLSSPFLPKLHPKFIWSPPLKSPHPSCLSISSVRIEDKPQTTTTGGREVAQPLKKETPPPSSPRKIRSSTLSTTRPSLEPFSFPSTIFNVVDNFINTF
VDPPLRPSVDPRYVLCDNFAPVGELPPTQCLLVEGSLPASLNGAYIRNGPNPQFLPRGPYHLFDGDGMLHSIRISDGKATLCSRYVKTYRYTVENERGYP
VFPNVFSGFNGLRASAARGAVTAARVLAGEFNPINGAGLANTSLALIAEKLYALGESDLPYEVKVAPDGDIITHGRHDFDGKLFMSMTAHPKVDLDTGEA
FAFRYGPMPPFLTFFRIDPDGTKQPDVPIFSMTSPSFMHDFAITKKYAIFPEIQIGMNPMEMMAGGAPVGANPGKVPRLGVIPRYAKDESEMKWFDVPGF
NIIHAINSWEEDDGDTIVLVAPNILSVEHTLERMDLIHASVEKVKIDLKTGMVSRSPISTRNLDFGAINPAYTGKKNKYVYAAIGDPMPKISGLVKLDVS
VSDSDRRDCIIASRLFGEGCFGGEPFFVAREPNNPNADEDDGYVVTYVHNEKNGESSFLVMDAKSPNLDTVAAVKLPRRVPYGFHGIFVRENDLNKL
>NatCCD4.6
MDTLSSSFLPKLHPKFILSPPLKSPHPTCLSISSVRIEDKPQITTTTTSGRREVARPLKKETPSPPSSPRKIRSSTLSTMRPSLEPFSFPSTIFNVVDNF
INTFVDPPLRPSVDPRYVLCDNFAPVGELPPTQCLVVEGSLPTSLNGAYIRNGPNPQFLPRGPYHLFDGDGMLHSIRILDGKATLCSRYVKTYRYNVENE
RGYPVFPNVISGFNGLTASAARGAVTAARVLAGQFNPTNGVGLANTSLALIGGKLYALGESDLPYEVKVAPDGDIITHGRHDFDGKLFMSMTAHPKMDLD
TGEAFAFRYGPMPQIVTFFRIDPDGTKQPDVPIFSMTSPSFMHDFAITKKYAIFPEIQIGMNPMEMMAGGAPVGANPGKVPRLGVIPRYAKDESEMKWFD
VPGFNIIHAINSWEADDGDTIVLVAPNILSVEHTLERMDLIHASVEKVKIDLKTGMVSRSPISTRNLDFGVINPAYTGKKNKY
>NatCCD4.7
MSSSIHTPNSLHDSSIRIDISSTDNAISRSESNTSLIFNALDEFINKFIDPPRRPSVNPRHVLSGNFAPVDELSPTECEILEGFLPPCLDGVYVRNGPNP
QFFPQGPHHLFDGDGMLHSIKISQGKATFCSRYVKTYRYMLEREMGSSIIPHVFSSFNGLVACLARGVVSAARMLTGQFTHVNGLGVASTSLAFFGGNLY
ALDESDLPYAIRMAQDGDIITLGRCDFNGKLVTNMTAHPKTDQDSGETFAFRYSFMRPYLSFFRFDNDGIKQPDVPIFSMASPSFVHDFAITKNYAIFPQ
IQLEVRPLNMVFGSGSPLRVDQSRNPRIGVIPRYAKDETEIKWFKVPGLNILHAINAWEENSGDTIVMVAPNILTLEHFIERMDLFHSLIEKVKIDLKTG
TVSRHPLSTRNLEFGVINPAYVGKKNKYVYAGVEDLMPKTAGVVKLDITSMKCRDCVVASRFYGHGCYGGEPLFVAKEPNNPNADEDDGYLVTYVHDENT
GNSRFLVMDAKSATLEIVTAVTLPQRVPYGFHGLFVREEEL
>NatCCD4.8
MDAISSHLLPKLSHHQKFTCFHMPKRKPSSSHLPNSLNVSSIRIDASSSSSSTSTSAATDDVTTATTSSAAHLLKKQAPSRLKINTSMIFDALDEFITKF
IDPPLRPSTDPRYVLSGNFAPVDELSPTECEVLDGSLPSCLDGVYIRNGPNPQFLPSGPYHSFDGDGMLHSIKISQGKATFCSRYVKTYRYMLEHEMGSK
VIPHVFSGFNSLMACVARGTVSAARMLTGQYTHTSGIGLANTSVAFFGGSLYALGESDLPYAIKVTQEGDIITLGRCDFNGKLVTNMTAHPKTDPDTGET
FAFRYSAIRPFLTFFRFDANGKKQDDVPIFSMTSPSFVHDFAITKNYAIFPEIQLGMRPLDVIFGLGPLIGMDWSKVPRLGVITRYAKDETETKWFNVPG
FNILHAINAWEENGGDTIVLVAPNILRLEHCIERMDLIHSSIEKVKIDLKTRVVSRHPLSTRNLEFGVINPTYLGMKNRYVYASIGDPMPKVSGVVKLDV
SLSGQDSHDCVVASRLYESGCYGGEPFFVAKEQNNPNLDEDDGYLVSYVHNENTKESTFLVMDAKSATLEIIAAVNLPQRVPYGFHGLFGKTYSMIESQN
NTSEINVEGDPTHSKGPNNESVVLARR
```

Percent Identity  Matrix - created by Clustal2.1

| | | | | | | | | |
|---|---|---|---|---|---|---|---|---|
| 1: NatCCD4.1 | 100.00 | 68.62 | 55.65 | 55.48 | 54.31 | 46.46 | 53.90 | 51.99 |
| 2: NatCCD4.2 | 68.62 | 100.00 | 59.83 | 59.66 | 59.03 | 51.50 | 57.25 | 56.55 |
| 3: NatCCD4.3 | 55.65 | 59.83 | 100.00 | 98.66 | 80.98 | 71.27 | 63.89 | 62.86 |
| 4: NatCCD4.4 | 55.48 | 59.66 | 98.66 | 100.00 | 82.32 | 71.27 | 63.89 | 62.52 |
| 5: NatCCD4.5 | 54.31 | 59.03 | 80.98 | 82.32 | 100.00 | 84.52 | 64.63 | 63.37 |
| 6: NatCCD4.6 | 46.46 | 51.50 | 71.27 | 71.27 | 84.52 | 100.00 | 57.58 | 56.64 |
| 7: NatCCD4.7 | 53.90 | 57.25 | 63.89 | 63.89 | 64.63 | 57.58 | 100.00 | 76.52 |
| 8: NatCCD4.8 | 51.99 | 56.55 | 62.86 | 62.52 | 63.37 | 56.64 | 76.52 | 100.00 |

Fig. S1. CCD4 sequences identified *in N. arbor-tristis*. A) Prediction of 8 amino acid sequences of CCD4 enzymes encoded by genes identified in the genome of *N. arbrt-tritis*. B) Percent identity matrix among the CCD4 enzymes created by Clustal2.1.

Fig. S2

**NatCCD4.1**

**Prediction: Plastid, Soluble**

| Localization | Plastid | Cytoplasm | Mitochondrion | Endoplasmic reticulum | Nucleus | Peroxisome | Extracellular | Lysosome/Vacuole | Golgi apparatus | Cell membrane |
|---|---|---|---|---|---|---|---|---|---|---|
| Likelihood | 0.9999 | 0.0001 | 0 | 0 | 0 | 0 | 0 | 0 | 0 | 0 |

| Type | Soluble | Membrane |
|---|---|---|
| Likelihood | 0.8686 | 0.1314 |

**NatCCD4.2**

**Prediction: Plastid, Soluble**

| Localization | Plastid | Cytoplasm | Mitochondrion | Nucleus | Endoplasmic reticulum | Extracellular | Peroxisome | Lysosome/Vacuole | Golgi apparatus | Cell membrane |
|---|---|---|---|---|---|---|---|---|---|---|
| Likelihood | 1 | 0 | 0 | 0 | 0 | 0 | 0 | 0 | 0 | 0 |

| Type | Soluble | Membrane |
|---|---|---|
| Likelihood | 0.9107 | 0.0893 |

**NatCCD4.3**

**Prediction: Plastid, Soluble**

| Localization | Plastid | Cytoplasm | Mitochondrion | Nucleus | Endoplasmic reticulum | Extracellular | Peroxisome | Lysosome/Vacuole | Golgi apparatus | Cell membrane |
|---|---|---|---|---|---|---|---|---|---|---|
| Likelihood | 1 | 0 | 0 | 0 | 0 | 0 | 0 | 0 | 0 | 0 |

| Type | Soluble | Membrane |
|---|---|---|
| Likelihood | 0.8869 | 0.1131 |

**NatCCD4.4**

**Prediction: Plastid, Soluble**

| Localization | Plastid | Cytoplasm | Mitochondrion | Nucleus | Endoplasmic reticulum | Extracellular | Peroxisome | Lysosome/Vacuole | Golgi apparatus | Cell membrane |
|---|---|---|---|---|---|---|---|---|---|---|
| Likelihood | 1 | 0 | 0 | 0 | 0 | 0 | 0 | 0 | 0 | 0 |

| Type | Soluble | Membrane |
|---|---|---|
| Likelihood | 0.8844 | 0.1156 |

**NatCCD4.5**

**Prediction: Plastid, Soluble**

| Localization | Plastid | Cytoplasm | Mitochondrion | Nucleus | Endoplasmic reticulum | Extracellular | Peroxisome | Lysosome/Vacuole | Golgi apparatus | Cell membrane |
|---|---|---|---|---|---|---|---|---|---|---|
| Likelihood | 1 | 0 | 0 | 0 | 0 | 0 | 0 | 0 | 0 | 0 |

| Type | Soluble | Membrane |
|---|---|---|
| Likelihood | 0.9003 | 0.0997 |

**NatCCD4.6**

**Prediction: Plastid, Soluble**

| Localization | Plastid | Cytoplasm | Mitochondrion | Nucleus | Endoplasmic reticulum | Extracellular | Peroxisome | Lysosome/Vacuole | Golgi apparatus | Cell membrane |
|---|---|---|---|---|---|---|---|---|---|---|
| Likelihood | 1 | 0 | 0 | 0 | 0 | 0 | 0 | 0 | 0 | 0 |

| Type | Soluble | Membrane |
|---|---|---|
| Likelihood | 0.905 | 0.095 |

**NatCCD4.7**

**Prediction: Plastid, Soluble**

| Localization | Plastid | Mitochondrion | Cytoplasm | Nucleus | Extracellular | Peroxisome | Endoplasmic reticulum | Lysosome/Vacuole | Golgi apparatus | Cell membrane |
|---|---|---|---|---|---|---|---|---|---|---|
| Likelihood | 0.9992 | 0.0004 | 0.0003 | 0 | 0 | 0 | 0 | 0 | 0 | 0 |

| Type | Soluble | Membrane |
|---|---|---|
| Likelihood | 0.8146 | 0.1854 |

**NatCCD4.8**

**Prediction: Plastid, Soluble**

| Localization | Plastid | Mitochondrion | Cytoplasm | Nucleus | Endoplasmic reticulum | Extracellular | Lysosome/Vacuole | Peroxisome | Golgi apparatus | Cell membrane |
|---|---|---|---|---|---|---|---|---|---|---|
| Likelihood | 1 | 0 | 0 | 0 | 0 | 0 | 0 | 0 | 0 | 0 |

| Type | Soluble | Membrane |
|---|---|---|
| Likelihood | 0.8953 | 0.1047 |

Fig. S2. Predicted in silico localization results of the different NatCCD4 enzymes using DeeoLoc (https://services.healthtech.dtu.dk/services/DeepLoc-1.0/).

Fig. S3

```
P-NatCCd4.2      tcgtcatttgcgaagaatgacccaagtaattaactgatattctggtagaatcttgaaaca     60
P-NatCCd4.1      ----------------------ca---agtaaaccgcaattctgtagtgattgtaatata     35
                                       **   * * *** *  ******     **  * * * *

P-NatCCd4.2      atgacggcaagtataatcgtcc--cgaataccagccaaagcacttatctccaaattttag     118
P-NatCCd4.1      --------cagttcaatggtttggctaagaccaactaaaacttatggattagaatcggcg     87
                          ***  *** **    * ** **** * *** *   *   *   ***    *

P-NatCCd4.2      ctcaattattttcatcttaattttcatgtcgtccgaagctagatgaatatt---------     169
P-NatCCd4.1      tttttttcctagaattgggtttctcctgtgtaaacagtgtagagaaataatgtggctaag     147
                  *   **  *   **     ** ** ***      *   ****  **** *

P-NatCCd4.2      attatct---------ttattataataataataataacgaca-----------ttgccga     209
P-NatCCd4.1      actagctaaaacttctggattagaatcggtgtgtaagcagtatagagaaataatgcctga     207
                 * ** **           **** ***     *   * *   *           *  * **

P-NatCCd4.2      ttctac-------cttagtatcttctttacagttaaaatacatcttcatatcccaagaat     262
P-NatCCd4.1      caaaaactttggattccgccatttctttcaattttggcaaattcatcatctcaatagcta     267
                     *         *  *    ******  * **     *  ** **** **   **

P-NatCCd4.2      aaatattttgatgttatctttatgatctgttctgtcggctgcatttgttttattcatagt     322
P-NatCCd4.1      aaagagggca-----ctc--------cttgaagttcggctgcatgttt-----ccttcta     309
                 *** *           **        **      ********** * *      * *

P-NatCCd4.2      tttggagatgctttaggtttgatagaaatacaaaatgtttgaaaatggaaacagaatact     382
P-NatCCd4.1      tttgggaccaaataataattcaaatgcctaataaatgttattatgttgagacttgagaga     369
                 *****       * *   ** * *    **  *******   *  * ** **   * *

P-NatCCd4.2      atagaagggggtatcctgcaactggtaat-atatacattagtgcaaagacagatttactt     441
P-NatCCd4.1      agaaaaaatgttttagaaccaatggttggtttggtttttatcagtaagatgacttgactc     429
                 * * **   * * *    * * ****     *     ***     ****    ** ***

P-NatCCd4.2      agttaaacgtagcaacaaaatcaccacccctaaaaatcatgttacatcaatacatcccac     501
P-NatCCd4.1      aataagaaaaaacatctattttctttaccattatatatgtcctcgatctattcatcccga     489
                 * * * *   * ** * *  *      ** * * *    *  *  *** ** ******

P-NatCCd4.2      caacaggacgggcttgtttggcacagaaatgctatcgagattttagtttctttttcgacg     561
P-NatCCd4.1      tt---cccgaccccccgtctggctatggaaacaaatcgggctttgagtttctagtccgacc     546
                             *  ** ****   * **    **** * *** *******  * ****

P-NatCCd4.2      cagcttagttga---atcggccgcatcagcttcatttggagttcgatttgatcctcaaga     618
P-NatCCd4.1      accaattatagtcagatgacattactccctgtaattggtagtcgggctccaaattcaga-     605
                      *  * *    **        **    * *** * ***  *  *  *   ***

P-NatCCd4.2      catactcactatgaagtattttggataaaatcta--aatacagccaaggttctaaaaaac     676
P-NatCCd4.1      -----gcccgatctgtttttatagaaaatatgttccccgactacccccgttttcaaaaaa     660
                       * * **    * ** * ** ** ** *      **  **   *** * *****

P-NatCCd4.2      gtaaggcatagaaaaacgtggaggtcgagcattgagcctgtac---ttgtttaagcgtac     733
P-NatCCd4.1      aatccattttttaaccttttgaaatattttataaaatgtttttttagtatttcggccagg     720
                         *   **    * **  *     **  *   * *      * ***  **

P-NatCCd4.2      taatgcataataaaagcctcatcaatgcgtactaaaacgcaagacacgatgaaacgtgaa     793
P-NatCCd4.1      agttgtcaagtt-tcgaccacccaatttgac-----------------cttttagctgcg     762
                    **   * *    * *    ****  *                    *   *  **

P-NatCCd4.2      gtccaagcttcaaacataagcgccccc--------taagtattg--cctagatgagattt     843
P-NatCCd4.1      cccctaacttataagaaaacaatttgttggttttagaactatttttattttatgtattta     822
                   ** * ***  ** * **                 ** ****     *  ***   **

P-NatCCd4.2      ttagaatactaaatacagtaacatgacaatattctggtttaacactgtctcatggaaaac     903
P-NatCCd4.1      tttattttcgaa--atattattttcaatatatcttcttataccatctaattatatctaga     880
                 **    * * **  * * **   * *  ****  *  * ** **     * **    *

P-NatCCd4.2      cctgg---------ctgtcaaacgagggtacactagtgatg--ccc-acttgggatcaag     951
P-NatCCd4.1      catgatttagttaaatgtttcaagttcgtgacgtggtgacctggcaaagcttcaactaag     940
                 * **           ***   * *   **    * ****     *  *  *   *  ***

P-NatCCd4.2      tacatagattatccacat--taggttatttgtccactaatatgctcttatttaacatctt     1009
P-NatCCd4.1      ccagtcgtttgtgtttttaaattttttcctcttaactaatttgctcttatttaaactct-     999
                     * * ** *     *      **   * *  ****** *************  ***

P-NatCCd4.2      ctgtctcccttcatttgtcagagtgtagataaatcctgcagaatttgtt-----------     1058
P-NatCCd4.1      taatctctcttcatttgtcacaaggtacatcaatctcctgaatcttgattatcaatttga     1059
                    **** ************ *  *** ** ****      *  *** *

P-NatCCd4.2      -gtcaagaaagatttgactatgggaacactt     1088
P-NatCCd4.1      cgcccaaaatgact--agcatgggaacactt     1088
                  * * * ** ** *  *  ************
```

Supplementary Fig. S3. Nucleotide sequence alignment of NatCCD4.1 and NatCCD4.2 promoters. The star codon is labelled in red. Conserved residues are depicted with an asterisk.

Fig. S4

```
NatCCD4.1        MVEENHETSILPEKAFSSRTTLQKLVQLIGKSTPR----KEV------VKKKKEPSLVAT   50
NatCCD4.2        MVSSVR-TEDKPQTTTTNGTAGREVAQPLKKETPPLPNPREILPSFLAPKKMESLLSPST   59
                 **.. : *.  *:.: :. *: :::.* : *.**     :*:       ** :.    :*

NatCCD4.1        ILNAYEDFICTFLDLPLPPSLDPKHVLSGNYAAVDELPPTPCEVVEGALPSCLDGAYLRN   110
NatCCD4.2        IFNVVDNFINTCIDPPLHPSTDPKYVFLNNYAPVDELPPTQCQVVEGTLPTCLNGAYIRN   119
                 *:*. ::** * :* ** ** ***:*: .*** ******* *:****:**:**:***:**

NatCCD4.1        GPNPQFIPRGPYHLFDGDGMLHVIKISQGKAVFCSRYVKTHKYIIEHKNGSPVIPSIFSS   170
NatCCD4.2        GPNPQFFPRGPYHTFEGDGMLHSIRIFNGKATFCSRYVKTYRYIVENKNGFPIFPNVFSG   179
                 ******:****** *:****** *:* :***.********::**:*:*** *::*.:**.

NatCCD4.1        FNGFSATMARLVPFVVRSLAGQFNPSTHGIGVANTSLALIGGDLFALAESDLPYKLKLTP   230
NatCCD4.2        FNGLTALATRGALAAARILAGQFNPA-NGTGPANTSLALVGGKLYALGESDLPYTVKVAE   238
                 ***::*  :*  .   ..* *******: :* * *******:**.*:**.******.:*::

NatCCD4.1        DGDVITQGRHTFCSKPVMSMTAHPKTDPNTGEAFGFAYNVFHPFLTYFRINREGVKQKDI   290
NatCCD4.2        DGDIITQGRHDFDGKLIMSMTAHPKVDPDTKEAFAFRYGAISPFLTFFRINPDGTKQPDV   298
                 ***:****** *  .* :*********.**:* ***.* *..: ****:**** :*.** *:

NatCCD4.1        PINSLTRSSFIHDFAVTKNYVVFPETQIVIDPMNIIRGKSPVGTDLAQVPRLGILPRYAE   350
NatCCD4.2        PIFSMKSPSLIHDFAITKKYAIFPEIQIGINPLKMMAGGSLIGTNPGKVPRLGVIPRYAK   358
                 ** *:.  *:*****:**:*.:*** ** *:*:::: * * :**: .:*****::****:

NatCCD4.1        NDAEMTWIDAPGLNMVHAFNAWEEDDGNRVVILASNAFEFESPSDPMDFLHLSVDRVEIN   410
NatCCD4.2        DESEMKWFDVPGFNILHAINSWEEDDGDTIVLVAPNILSVEHTLERVDLIHLSVEKVKID   418
                 :::**.*:*.**:*::**:*:******: :*::* * :..*    : :*::****::*:*:

NatCCD4.1        LKEKKVLRQPLCSTNLDLAVINPSYAGKKNRFIYAAVTTTTPKRAGVMKLDLSLFEADNR   470
NatCCD4.2        LKTGMVSRIPLSTRNLDFGAINPSYTGKKNKYVYAAIGDPMPKIPGVVKLDISVSGSDHR   478
                 **    * * **.: ***:..*****:****:::***:    **  **:***:*:   :*:*

NatCCD4.1        DCAVGSRLYEPGCFGGEPFFVPRDPGSREDVDEDDGYLVTYLHNEKTGESRFLVMDAKSP   530
NatCCD4.2        DCIVASRLFGEGCLGGEPFFVAKEPDN-PSADEDDGYVVTYVHNEKNGESSFLVMDAKSP   537
                 ** *.***:  **:******* ::*..  ..******:***:****.*** *********

NatCCD4.1        ELEIVAQVKLPGRVPYGFHGLFVPESELKDM   561
NatCCD4.2        NLDTVAAVKLPRRVPYGFHGIFVRENDLNKL   568
                 :*: ** **** ********:** *.:*:.:
```

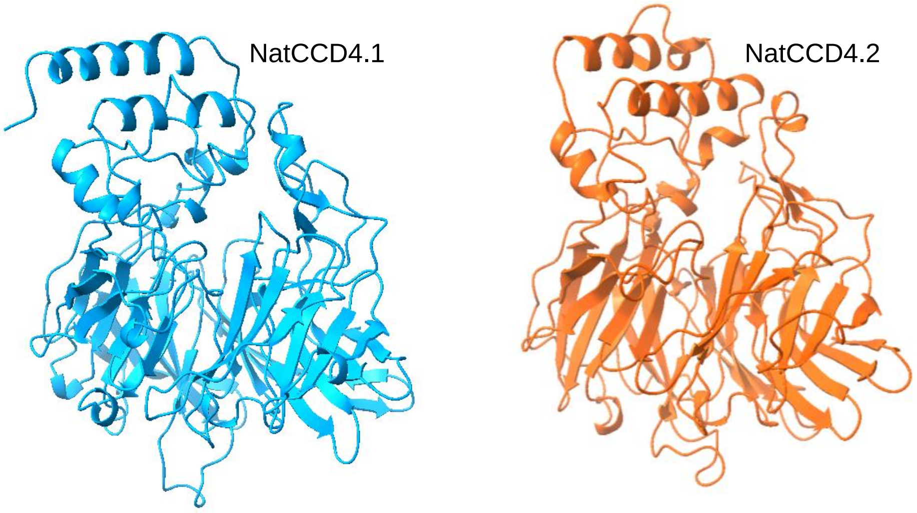


Fig. S4. Comparison of NatCCD4.1 and NatCCD4.2. A) Amino acid sequence alignment of NatCCD4.1 and NatCCD4.2. Conserved amino acid residues are depicted with asterisk, while two or one dost indicates conservation between groups of strongly or weakly similar properties, respectively. The amino acid residues involved in iron coordination are highlighted in yellow and blue, in yellow the conserved His residues. Tridimensional structures. The 3D structure were predicted using Phyre2 software at intensive mode (http://www.sbg.bio.ic.ac.uk/phyre2/ ) and Chimera X (https://www.cgl.ucsf.edu/chimerax/).

Fig. S5

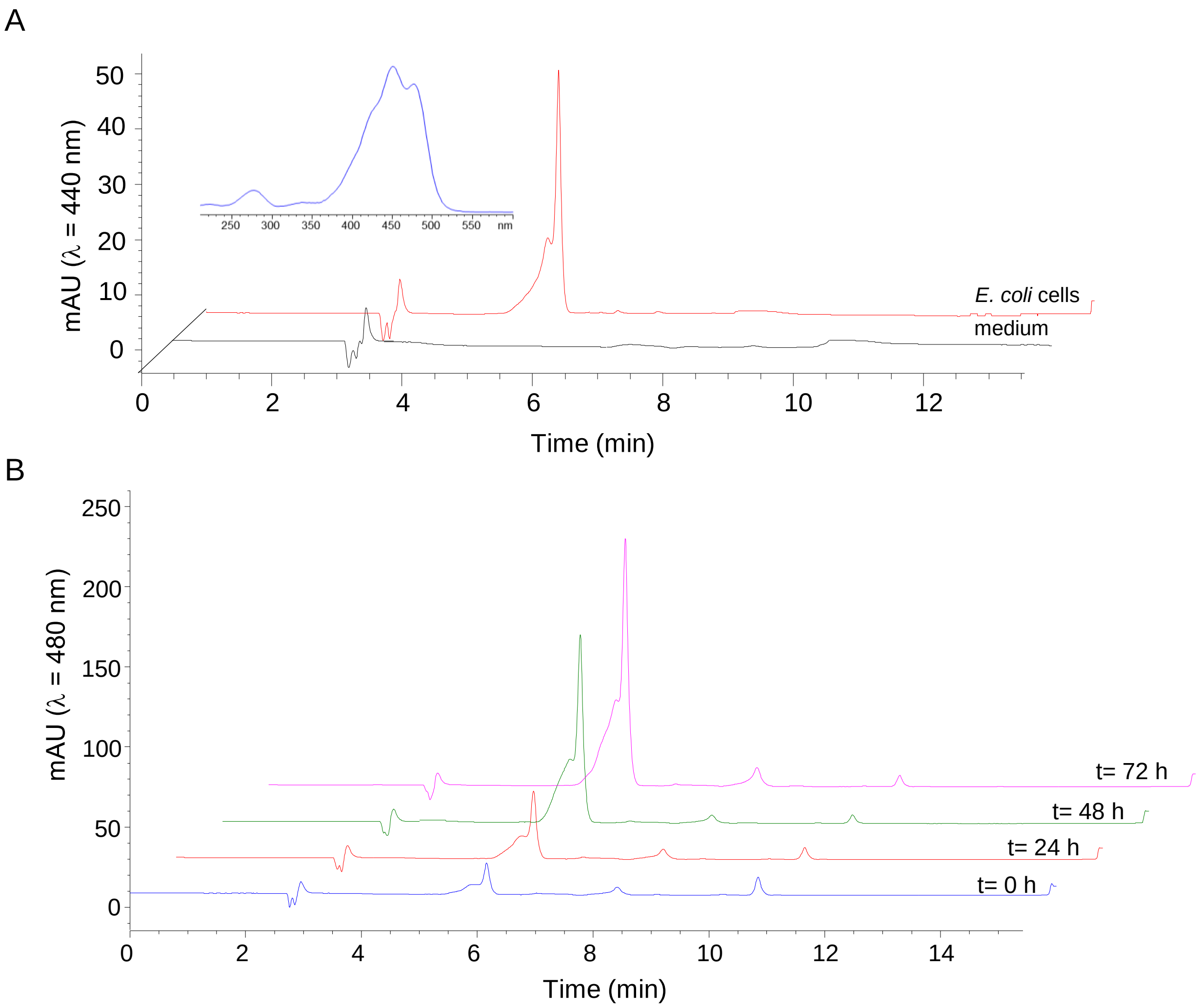


Supplementary Fig. S5. Lack of crocetin dialdehyde in cells pellet and medium in the two-phase experiment with n-dodecane. A) HPLC-DAD profile of apolar extracts from pellet of *E. coli* cells producing zeaxanthin and expressing NatCCD4.1 induced for 48 h with arabinose (0.2%, w/v) and with the addition of n-dodecane (16% v/v); and .HPLC-DAD profile of the medium in which these cells were grown. Inset is shown the spectra of zeaxanthin. B) HPLC-DAD profiles of apolar extracts from pellet of *E. coli* cells producing zeaxanthin and expressing NatCCD4.1 induced with arabinose (0.2%, w/v) and with the addition of n-dodecane (16% v/v) and collected at different times after induction.

Fig. S6

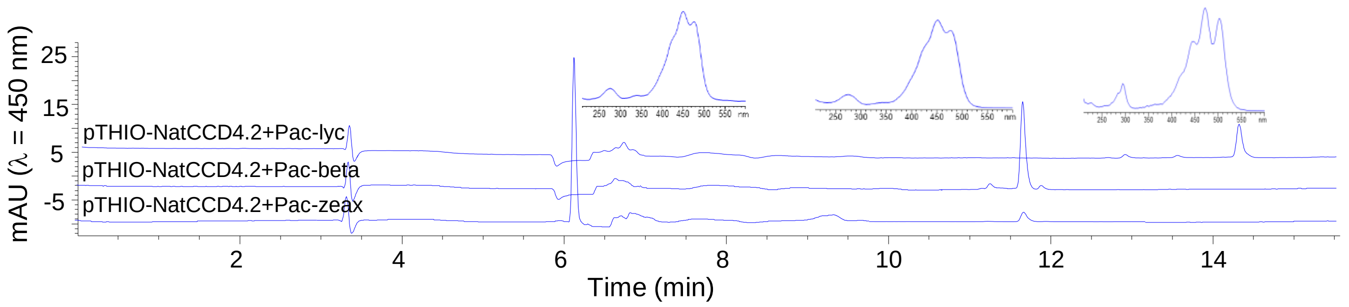


Supplemental Fig. S6. Crocetin dialdehyde was not detected in the n-dodecane phase collected from bacterial cells acumulating different carotenoids and expressing NatCCD4.2. The induction of NatCCD4.2 was promoted by the addition of arabinose (0.2% w/v), and the n-dodecane pase collected 24 h after induction. The experimentes were repeated independently three times with the same negative results,

Fig. S7

A

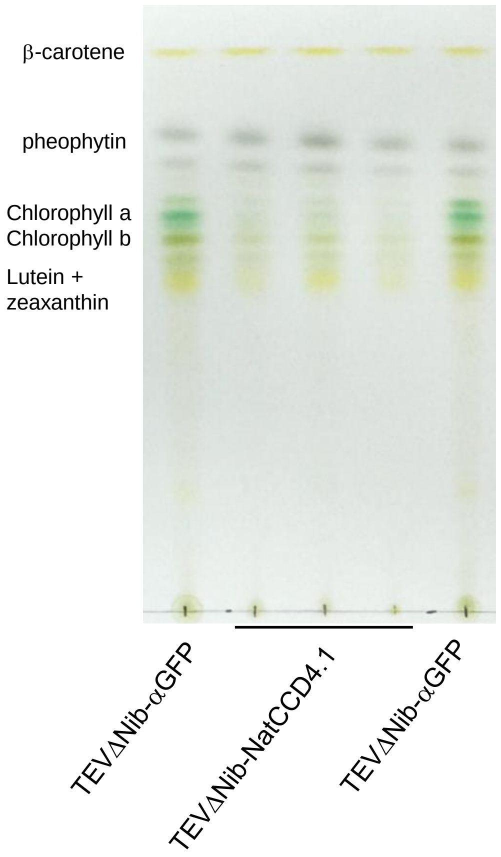


B

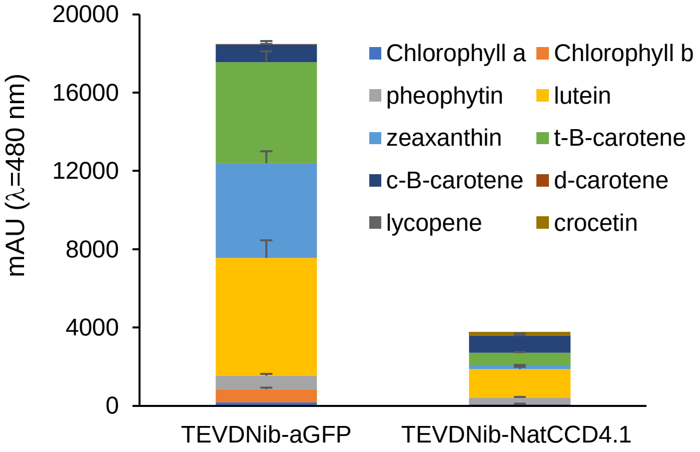


C

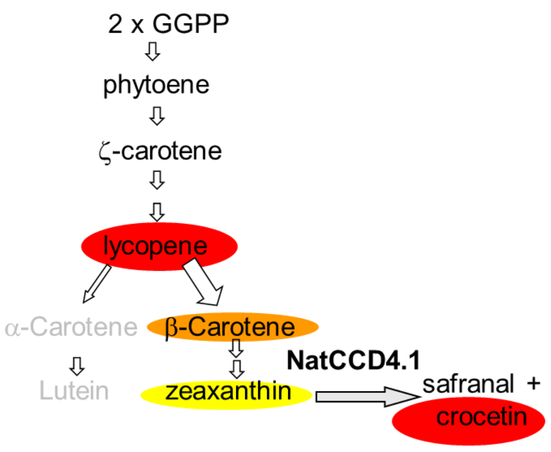


Supplemental Fig. S7. Analyses of apolar extracts from *N. benthamina* experiments. A) TLC analyses of pigments extracted from *N. benthamina* leaves with 2:1 methanol:chloroform. B) Levesl of carotenoids, chlorophylls and crocetin in the apolar extracts obtained from *N. benthamina* leaves . Data are averages ± SD of three biological replicates. Analyses were performed at 14 dpi. C) Schematic representation of the carotenoid pathway showing how NatCCD4.1 activity reduced notably the levels of the substrate used, zeaxanthin, and the levesl of the zeaxanthin precursor : β-carotene.

Fig. S8

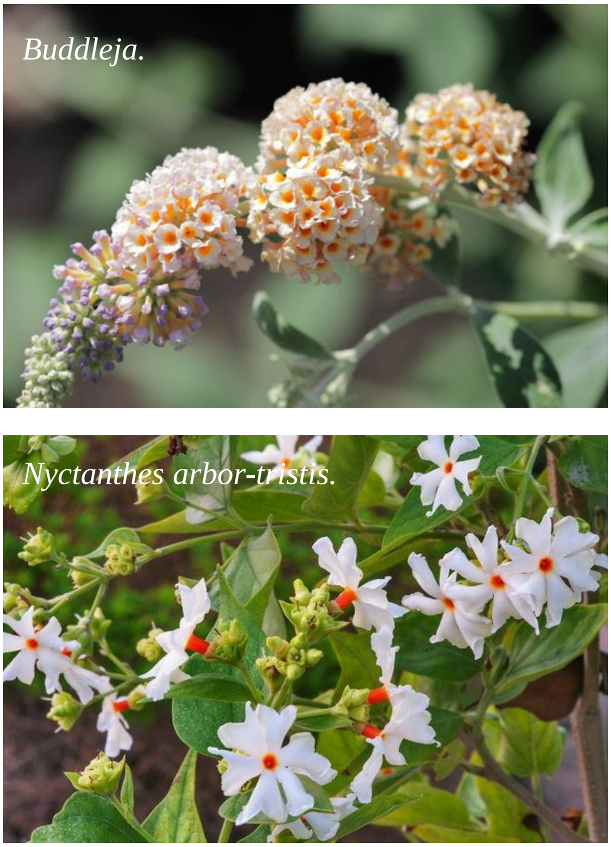


Supplemental Fig. S8. Crocins accumulated in the calisxof the flowers of *Buddleja* and *Nyctanthes arbor-tristis.*
